# Mitigation of Reproductive Function in Sodium Fluoride-administered Wistar Rats by *Tamarindus indica* Fruit Pulp Ethanol Extract

**DOI:** 10.1101/2024.10.01.615988

**Authors:** Lutfat A. Usman, Emmanuel O. Ajani, Muhammed A. Ishiaku, Taofeeq A. Bankole, Rasheed B. Ibrahim, Dauda K. Saka

## Abstract

Infertility is a stigmatized and emotionally challenging issue for affected individuals, often compounded by the high costs of conventional fertility treatments. This study investigated the prophylactic and therapeutic effects of an ethanolic extract of Tamarindus indica fruit pulp on sodium fluoride-induced sexual and reproductive dysfunction in Wistar rats. The extract was prepared via ethanol maceration, followed by filtration and drying. Seventy Wistar rats were used, with an initial acute toxicity study indicating no adverse effects at 5000 mg/kg of the extract.

Following toxicity assessment, rats were divided into seven groups, with varying treatments involving Tamarindus indica extract, sodium fluoride, and clomid. After the treatment period, biochemical analyses revealed that rats treated with 400 mg/kg of Tamarindus indica extract showed significant improvements (p < 0.05) in sperm count, luteinizing hormone (in males), testosterone, estrogen, along with reduced levels of follicle-stimulating hormone (in males), and luteinizing hormone (in females). Histopathological examination revealed normal histomorphology in certain groups treated with the extract. Phytochemical analysis identified the presence of tannins and steroids in the extract.

The study concluded that Tamarindus indica fruit pulp extract exhibits both prophylactic and therapeutic effects on sodium fluoride-induced sexual and reproductive dysfunction in rats, with the most significant effects observed at a 400 mg/kg dosage. These findings suggest that Tamarindus indica extract could serve as an alternative to conventional treatments for infertility.

## INTRODUCTION

In 2010, an estimated 48.5 million couples worldwide were infertile (Mascarenhas *et al*., 2012). Most of infertile couples have one of three major causes including a male factor, ovulatory dysfunction, or tubal-peritoneal disease (Kakarla and Bradshaw, 2008). Although several management practices and fertility techniques exist, they are however expensive, and beyond the reach of the poor. Moreover, they are not 100% accurate. Thus, there is a need to source for cheaper and more effective alternatives.

In developing countries, traditional medicine particularly medicinal plants, thanks to their accessibility, availability, and affordability are generally the first recourse of infertile couples (WHO, 2002). The use of medicinal plants in the treatment of diseases and goes back to several millennia and has considerably contributed to the development of pharmaceuticals. Reports indicates that up to 60% of the world’s population uses herbal products for medical purposes (Rates, 2001). Since two decades, the evaluation of natural materials as a source of potential drugs has been of resurgent interest in developing countries as well as in the developed ones. This growing interest for phytotherapy is due to several reasons namely; conventional medicine can be inefficient (ineffective therapy), abusive and/or incorrect use of synthetic drugs results in side effects and other problems, finding of the “natural”, large therapeutic spectrum of plant products and their effectiveness in the treatment of chronic diseases, need for development of new drugs etc. Several extracts, fractions or molecules isolated from these plants are today largely used to treat or relieve different aspects of infertility such as: absence of libido, sexual asthenia, erectile dysfunction, ejaculatory and relaxation dysfunctions, loss of orgasm, sperm abnormalities, ovarian disorders, menstrual irregularities, tubal peritoneal disease and fallopian tube blockage.

Over several years, researches have been conducted on the fertility efficacy of some ethnomedicinal plants. However, the toxicity, pharmacology and mechanism of action of these medicinal plants remains unclear. In view of the facts that medicinal plants have been an important ingredient in oriental herbal medicine for the prevention and management of several ailments and coupled with the fact that the traditional use of tamarind among some local people in Nigeria in managing infertility has been widely reported, it is therefore expedient to evaluate the effect of *Tamarindus indica* fruit pulp on the reproductive function.

Tamarind (*Tamarindus indica*) is a leguminous tree in the family Fabaceae that is indigenous to tropical Africa. The tamarind tree produces edible, pod-like fruits which are used extensively in cuisines around the world. The fruits are reported to have hypo-lipidemic, anti-inflammatory, anti-fungal and anti-bacterial properties (John *et al*., 2004). Tamarind pulp typically contains 20.6% water, 3.1% protein, 0.4% fat, 70.8% carbohydrates, 3.0 % fibre and 2.1 % ash (El Siddig *et al*., 1999). Virtually every part of *Tamarindus indica* (wood, root, leaves, bark and fruits) has either nutritional or medicinal value, with a number of industrial and commercial applications (Emmy *et al*., 2007). Tamarind is useful traditionally in gastric disorders, bilious vomiting, scurvy, datura poisoning, alcoholic intoxication, scabies, otalgia, stomatitis, constipation, haemorrhoids and eye diseases. Tamarind pulp is also said to aid in the cure of malarial fever (Timyan, 1996; Agnivesha *et al*., 2008). The leaves have a proven hepato-protective activity associated with the presence of poly-hydroxylated compounds, with many of them flavonolic in nature (El-Siddig *et al*., 2004). Due to their antimicrobial, antifungal, and antiseptic effects; tamarind leaves have extensive ethnobotanical use. Although a lot of studies have reported on the medicinal use of tamarind leaves, very few studies have reported on the medicinal use of the fruit pulp. Moreover, no reproductive model study has been reported yet in the literature to evaluate the efficacy of *Tamarindus indica* L. fruit pulp extracts on reproductive dysfunction in a living system. Therefore, the present study aimed to evaluate the efficacy of ethanolic extract of *Tamarindus indica* fruit pulp in the treatment of reproductive dysfunction in rats and compare the efficacy with Clomid. This is with a view to the development of a safer and cost-effective therapy for managing human sexual dysfunction.

## MATERIALS AND METHODS

### Drug Sample and chemicals

Clomifene citrate (clomid) was a product of *Bruno Farmaceutici* S.p.A and was purchased at Farason Pharmaceuticals Ilorin, Kwara State. Other chemicals and kits used in this study were obtained from Merck (India) and Sigma-Aldrich (Saint Louis, Missouri).

### Plant material

#### Collection and Authentication

The compact fruit of *Tamarindus indica* were obtained from farm land around Kwara State University, Malete campus in the month of September, 2018. The plant material was identified and authenticated by a botanist at the Herbarium of the Department of Plant Biology, University of Ilorin, Nigeria, where a voucher specimen with number UILH/001/1155 for *Tamarindus indica* was deposited.

#### Extraction

Dust free clean pulps were dried in an oven at 45^0^C. The dried fruit (916.77g) was then macerated in 2338 ml of 70% ethanol for 72 hours. The resulting macerate was blended to smoothness using a high-speed electric blender. Afterwards, the blended macerate was filtered using a muslin cloth, cotton-wool plug and Whatman No. 1 filter paper respectively. The resulting filtrate was concentrated to dryness using a vacuum oven at 60°C until a constant weight was achieved. The crude extract was stored in an air tight container and was placed in the refrigerator at a temperature of 4ºC prior to use.

### Experimental Protocols

#### Experimental Animals

Seventy clinically healthy rats of both sexes (in same proportion) with an average weight of 90 ± 5 g were obtained from Umaraahmarkeen Nigeria Global Ventures, Kwara State, Nigeria. The animals were maintained in stainless steel cages at the animal holding unit of the Department of Medical Biochemistry and Pharmacology, Kwara State University, Malete with optimum environmental conditions (temperature: 23 ±1°C, photoperiod: 12 h light/dark cycle, relative humidity: 45-50%). The animals were allowed free access to food and fresh water *ad libitum*. All the animals were acclimatized to laboratory conditions for a week before commencement of the experiment. The study was executed following approval from the Ethical Committee on the use of Laboratory Animals of the Department of Biochemistry, Kwara State University, Malete, Nigeria. Animal management and maintenance was per the principles of laboratory animal care (NIH publication no. 82-23, revised 1996) guidelines.

#### Acute Toxicity Test

This was carried out according to the World Health Organization (WHO) and the Organization of Economic Co-operation and Development (OECD) guidelines for testing of chemicals (Hashemi *et al*., 2008). Standard two phase approach (Lorke, 1983) was used in this approach. In the first phase, 9 rats were divided at random into three groups of three rats each. Food and water were withdrawn from the rats 12 hours before the experiment. Group I, II and III were orally administered with *E. chlorantha* extract at 10 mg/kg, 100 mg/kg and 1000mg/kg body weight respectively. Rats were observed for 48 hours for signs of toxicity or mortality. In the second phase of the experiment, three animals were assigned into three groups (IV, V and VI) of one animal each and thereafter administered with 1600mg/kg, 2900mg/kg and 5000mg/kg body weight of the extract respectively. They were then monitored for death over a period of 48 hours. LD_50_ was then calculated as

LD_50_ ≥ Maximum dose – Y**/** number of rats per group

Where Y = Sum of mean death

### Therapeutic study

#### Induction of Infertility

Experimental rats were rendered infertile after 45 days of continuous oral administration of 20 mg/kg of sodium fluoride (NaF) daily (Zhou *et al*., 2013; Zhang *et al*., 2016).

#### Animal grouping and Treatments

Seventy (70) rats were randomized into seven groups of 10 rats (5 males and 5 females) per group. The rats were labelled and treated as follow: Control (NC): non induced and administered with normal saline; Model group(MG): administered with NaF and normal saline; Preventive Group 1(PG1): first administered with 200 mg/kg b.w of *Tamarindus indica* fruit extract for 4 weeks followed by concurrent administration of NaF for another 45 days; Preventive Group 2 (PG2): first administered with 400 mg/kg b.w of *Tamarind indica* fruit extract for 4 weeks followed by the concurrent administration of NaF for the next 45 days; Treatment Group 1 (TG1): initially administered with NaF for the first 45 days after which they were concurrently treated with 200 mg/kg b.w of *Tamarindus indica* fruit extract for another 4 weeks; Treatment Group 2 (TG2): initially administered with NaF for 45 days and thereafter simultaneously treated with 400 mg/kg b.w of *Tamarindus indica* fruit extract for the next 4 weeks; Positive Control (PC): first administered with NaF for 45 days and then simultaneously treated with Clomid for another 4 weeks. NAF was administered at 20 mg/kg and clomid at 50mg/kg. All treatments were administered orally as a single dose per day using an oral intubator every day. Twenty-four hours after the last treatment, under mild diethylether anesthesia, the animals were sacrificed and blood was obtained from the jugular vein. Blood sample was transferred into plain centrifuge tube and allowed to clot at room temperature. It was then centrifuged within 1 hour of collection at 4000x g for 10 minutes to obtain the serum.

#### Preparation of Homogenates

The testes were harvested for histopathological analyses. The isolated testes were blotted with clean tissue paper and cleaned of fat. The testes were immediately fixed in Bouin’s fluid. These were then used for histological examination. The cauda epididymis was chopped, collected and placed in saline solution (0.9% NaCl, 5 mL) immediately. The fluid was incubated in a water bath at 37^0^C for 5 minutes to allow sperm to leave the epididymal tubules.

#### Histopathological Analysis

The method of Drury and Wallington (1980) was adapted for the histopathological examination of the harvested testes sections. Microscopic features of the cells of the treated rats were compared with both control and the model groups.

### Biochemical Assay Procedure

#### Sperm parameters study

##### Sperm count

Sperm count was determined using Mallassez hemocytometer. The sperm count was expressed as the number of sperms per milliliter of solution.

##### Sperm motility

The fluid from the cauda epididymis was obtained with a pipette and diluted with Tris buffer solution (2 mL). Immediately after their isolation, the sperm motility was evaluated microscopically at 400× magnification. The sperm forward motility was expressed as a percentage of motile sperms to total sperms counted.

##### Sperm viability

The ratio of live sperms to dead ones was evaluated using 1% trypan blue staining. The result was expressed as a percentage of the live sperms.

### Hormonal Assay

The levels of serum progesterone, testosterone, estrogen, follicle-stimulating hormone (FSH) and luteinizing hormone (LH) were determined by immunoenzymometric assay in the plasma of both male and female rats accordingly using Life Span BioSciences, Inc (LSBio) assay kits. Assays were carried out as described by the manufacturer.

### Statistical Analysis

All data were expressed as the mean of five replicates ± standard error of mean (S.E.M). Statistical evaluation of data was performed using SPSS version 16.0 using one-way analysis of variance (ANOVA), followed by Duncan’s posthoc test for multiple comparisons. Values were considered statistically significant at *p* < 0.05 (confidence level = 95%).

## RESULTS

### Acute Toxicity

There was no observable adverse effect in rats administered with 1000, 2000, 3000, 4000 and 5000 mg/kg body weight of the ethanolic *Tamarindus indica* fruit extract.

### Hormonal Assay

The result of the follicle stimulating hormone of the male rats (Figure 1) showed a significant reduction in the serum FSH level of the model group. Pre-treatment with *Tamarindus indica* extract raised the FSH level above that of the model group. The FSH concentration of the 400 mg/kg pre-treated rats were not different from that of the control value whereas the value recorded for the rats pre-treated with 200 mg/kg was observed to be significantly (p<0.05) lower than the control value. The result also showed that when the animals were treated with the extract (at the two tested dosage) after the induction of reproductive dysfunction, the observed FSH concentrations were not significantly different from that of the control group. Similar results were obtained for FSH level of the female rats. It was however observed that although the observed FSH level in the female rats treated with 200 mg/kg dose of the extract was lower than that of the model group, the observed value was higher than that of the control group (*p*<0.05). When compared between the gender, the FSH level of the male control group was significantly higher than that of the female group whereas the FSH level of the female model group was observed to be significantly higher than that of the male counterpart.

**FIGURE 1:**
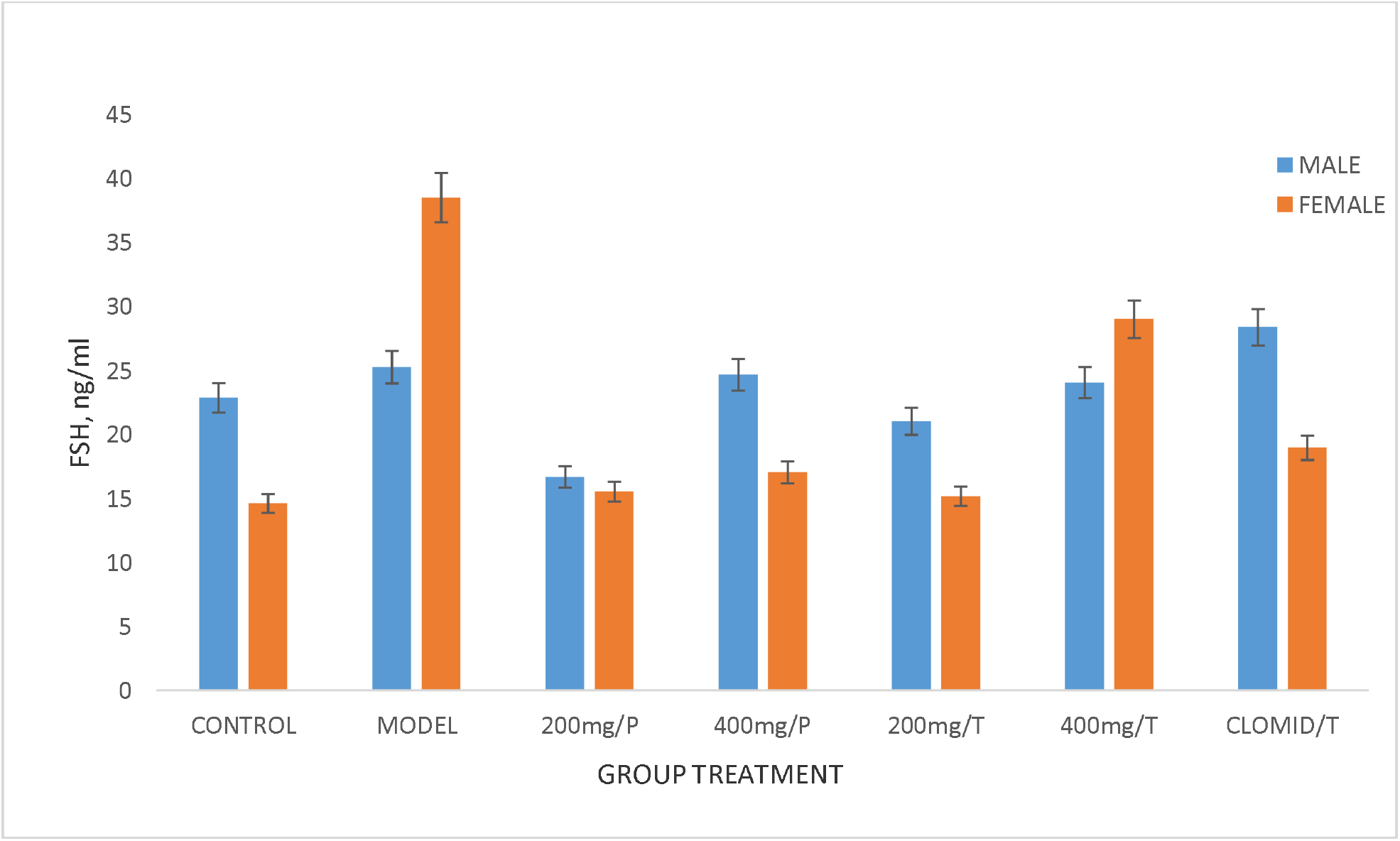
Effect of *Tamarindus indica* fruit pulp extract on serum follicle stimulating hormone (FSH) levels in Wistar rats. Results are Mean ±S.E.M, (n=5).

Figure 2 is the result on the effect of treatments on serum LH. A significant (p<0.05) reduction in the LH was observed in the model male rats when compared with the control group whereas, the model female rats group showed a significant elevation in the LH concentration when compared with the control rats. Pre-treatment with the extract in the male rats (at both doses) and post-treatment (at both doses) after NAF administration significantly raised the LH concentrations above the control group. However, no significant difference (p>0.05) was observed when the LH was compared among all the treated male groups. The LH concentrations of the female rats treated with the extract both prior to and after NAF administration (except in the group pre-treated with 200 mg/kg dose) was observed to be significantly lower than that of the model group. The result also showed that pre-treatment with the extract at 400 mg/kg reduce the LH of the female rats to the pre-treatment level.

**FIGURE 2:**
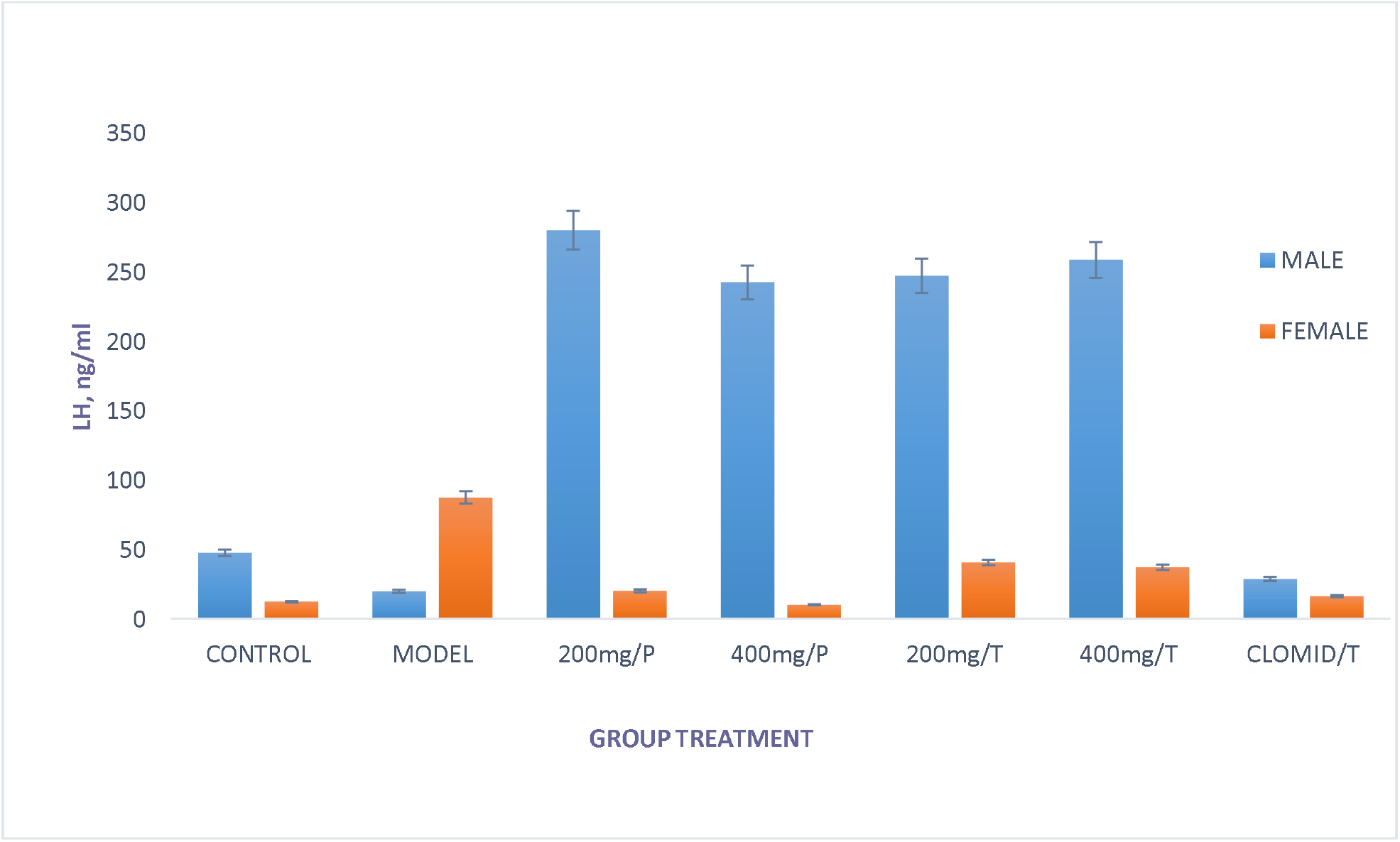
Effect of *Tamarindus indica* fruit pulp extract on serum leutinizing hormone (LH) levels in Wistar rats. Results are Mean ±S.E.M, (n=5).

The result of the effect of treatment on the testosterone concentration showed a significant reduction in the testosterone level of the model male rats when compared with the observed level in the control group (Figure 3). Treatment with the extract (in a dose dependent manner) and pre-treatment at 400 mg/kg and with clomid raised the serum testosterone level above the control value

**FIGURE 3:**
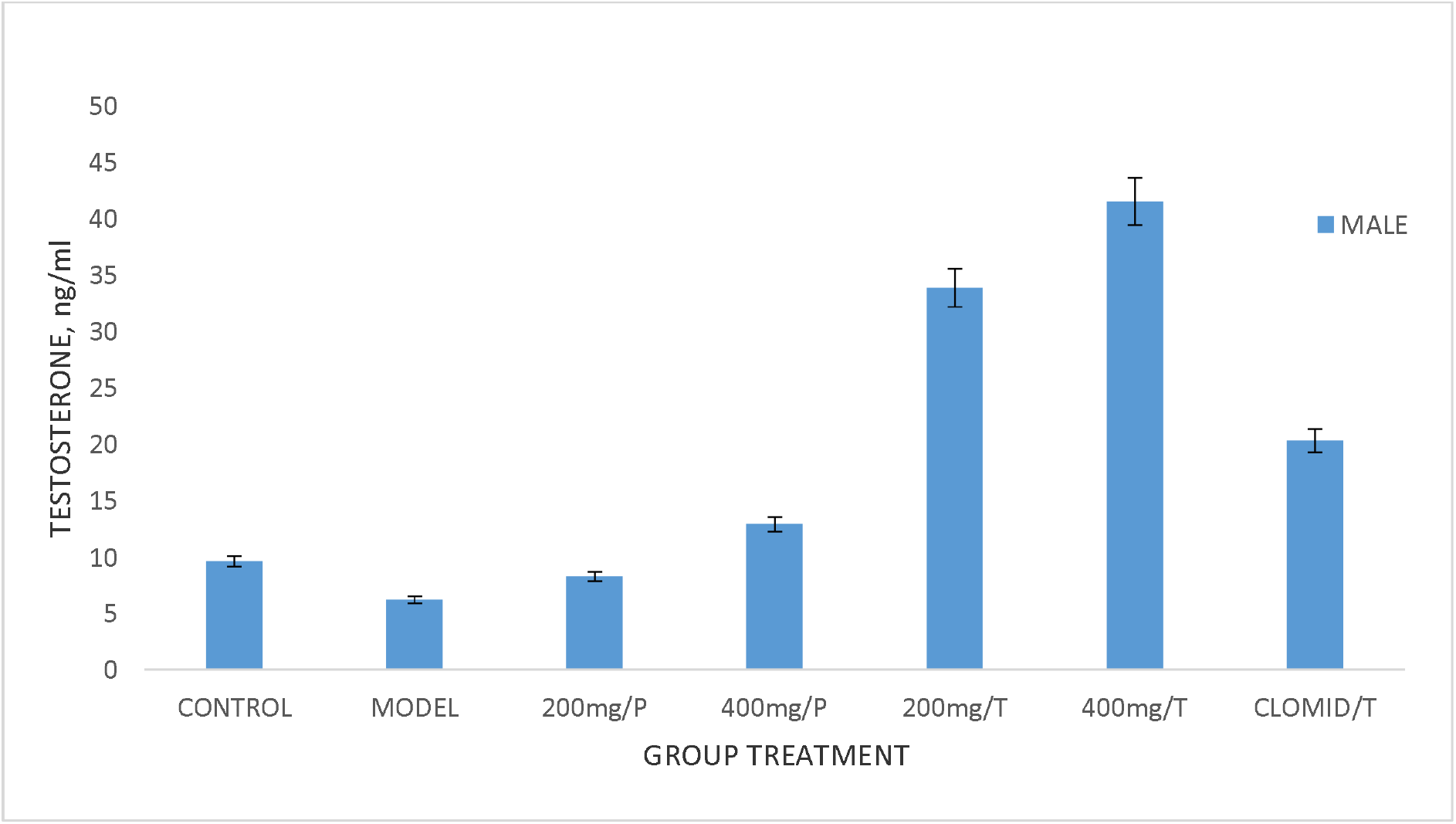
Effect of *Tamarindus indica* fruit pulp extract on serum testosterone levels in male Wistar rats. Results are Mean ±S.E.M, (n=5).

Figure 4 is the result of the effect of treatment on the serum oestrogen level of the female rats after treatment. Administration of NAF was observed to lower the serum oestrogen level of the model group below that of the control value. Treatment with the extract at 400 mg/kg dose was observed in the study to raise the serum oestrogen level above that of the control concentration. The serum oestrogen concentrations of all other treated groups were significantly not different (p>0.05) from that of the model group and lower than the control value.

**FIGURE 4:**
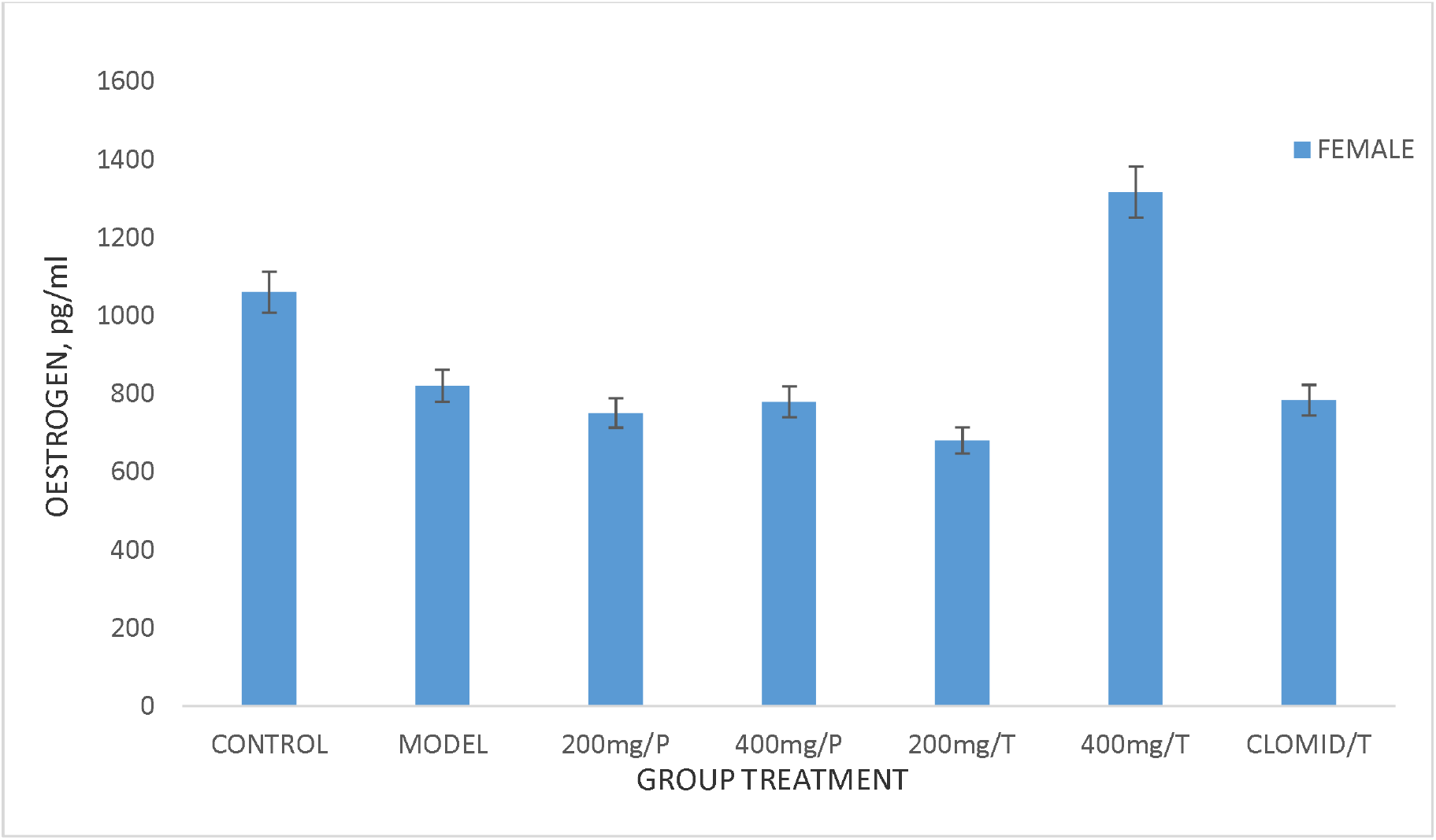
Effect of *Tamarindus indica* fruit pulp extract on serum oestrogen levels in female Wistar rats. Results are Mean ±S.E.M, (n=5). Bars with different letters are significantly different (*p*<0.05).

The result of the effect of treatment on serum progesterone concentrations in all experimental groups is shown in Figure 5. The model group showed a progesterone concentration that was significantly higher than that of the control value. Treatment with the extract prior to and after NAF administrations (at both doses) significantly lowered the progesterone concentration below that of the model group although the values obtained were significantly higher than that of the control group. The observed progesterone concentration in the rats treated with 400 mg/kg dose of the extract was not significantly different from that of the clomid treated rats.

**FIGURE 5:**
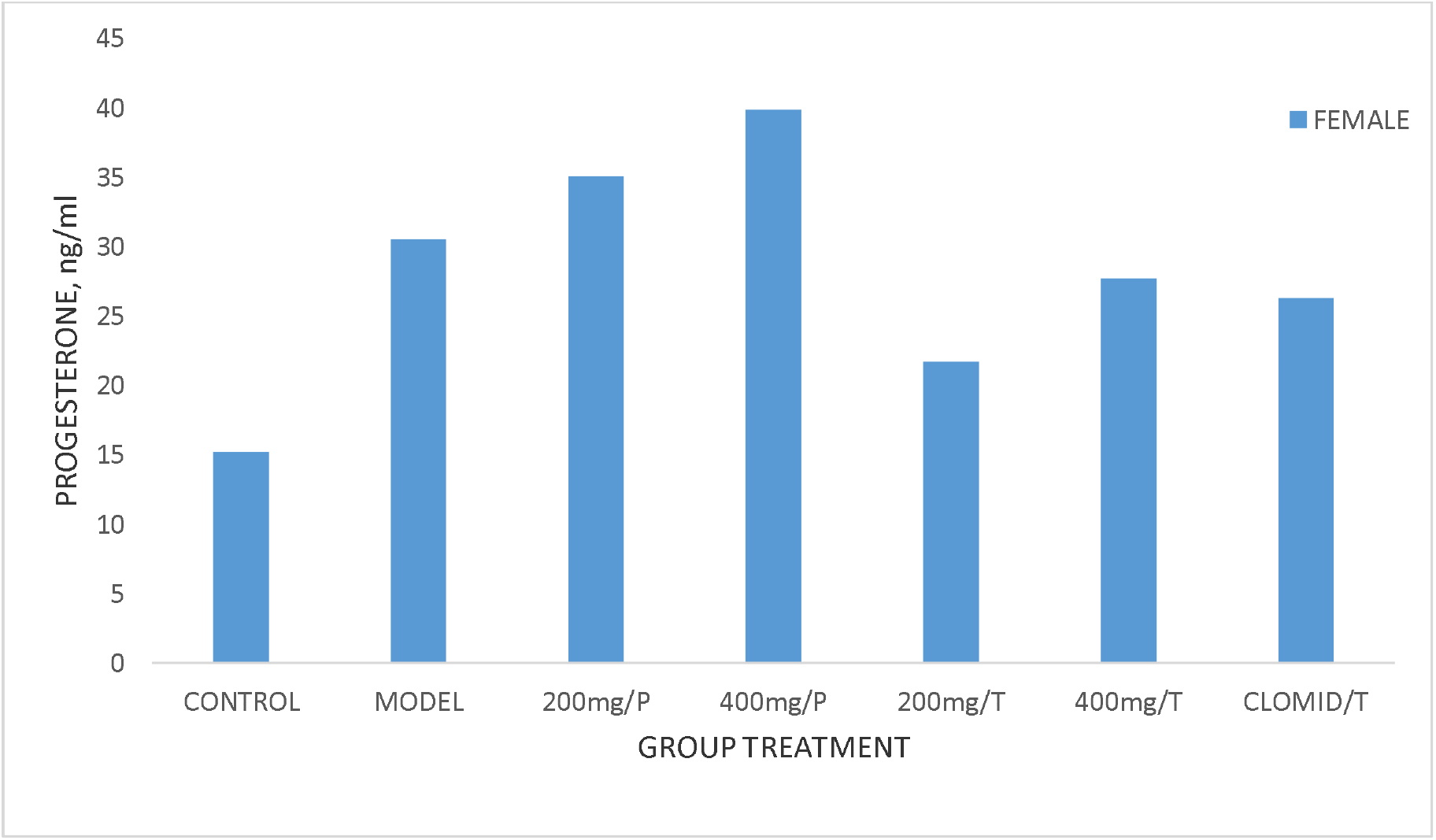
Effect of *Tamarindus indica* fruit pulp extract on serum progesterone levels in female Wistar rats. Results are Mean ±S.E.M, (n=5). Bars with different letters are significantly different (*p*<0.05).

### Sperm analysis

Figure 6 shows the result of the sperm count after treatment. No sperm count was recorded in the model group after NAF administration. Pre-treatment with the extract (at both doses) and treatment after NAF administration significantly raised the sperm count of the rats above the control value; this can be due to the sterols identified in the *Tamarindus indica* fruit extract. The highest sperm count value was observed in the rats treated with 400 mg/kg dose of the extract.

**FIGURE 6:**
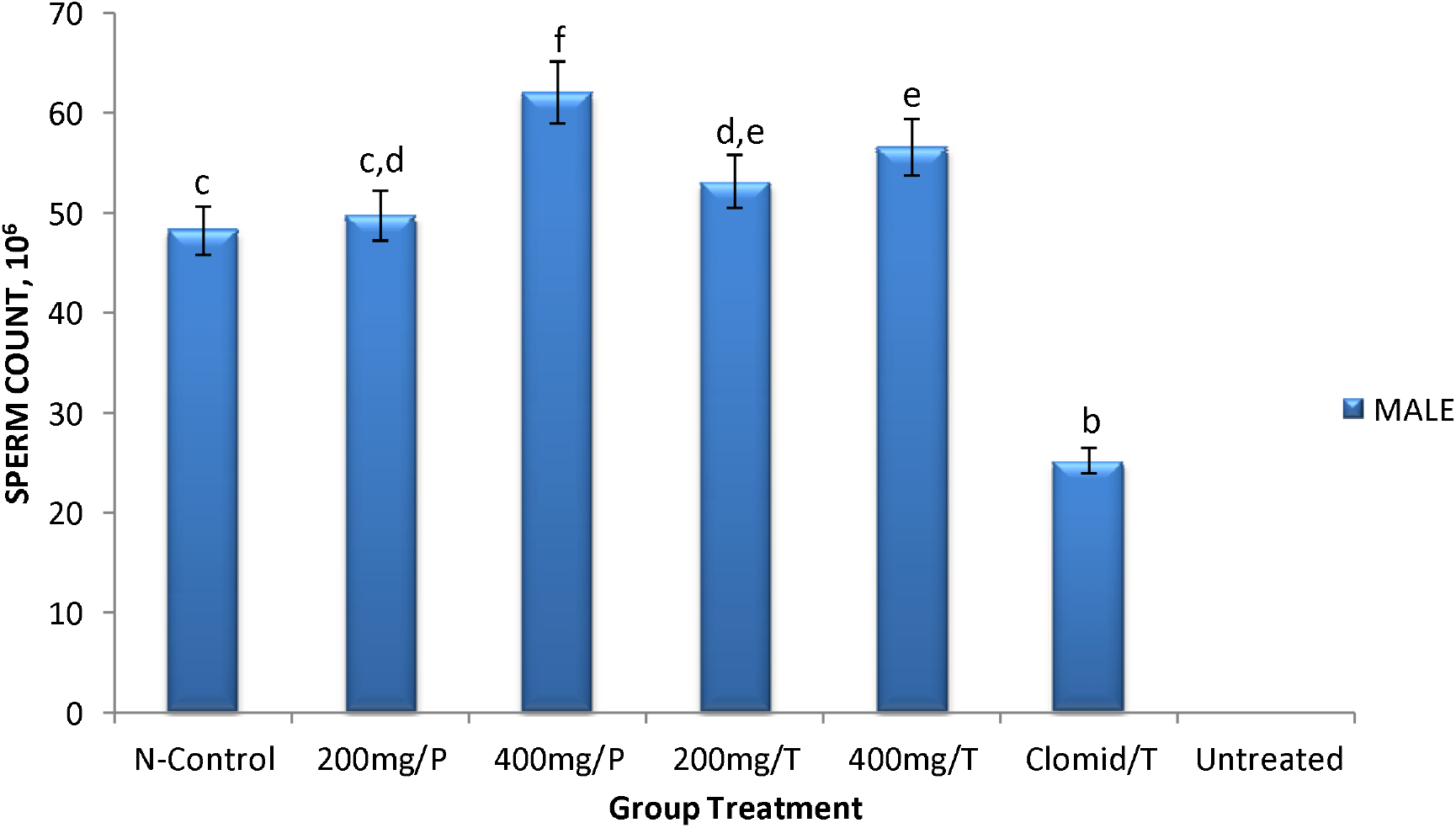
Effect of *Tamarindus indica* fruit pulp extract on sperm count in male Wistar rats with sodium fluoride induced reproductive dysfunction. Results are Mean ±S.E.M, (n=5). Bar with different letters are significantly different (*p* 0.05).

The sperm motility result also showed no motile sperm in the model group after treatment (Figure 7). The motility of the sperm recorded in the treatment groups were not statistical different from the control group. Similarly, no sperm morphology was observed in the model group (Figure 8) after NAF treatment. Although sperm morphology was observed in all other groups, the observed values in all the treatment groups were however significantly lower than that of the control value. The sperm viability result also showed that no viable sperm was observed in the model group. Although all other treatment groups showed viable sperm, the observed values were not significantly different from the control value (Figure 8)

**FIGURE 7:**
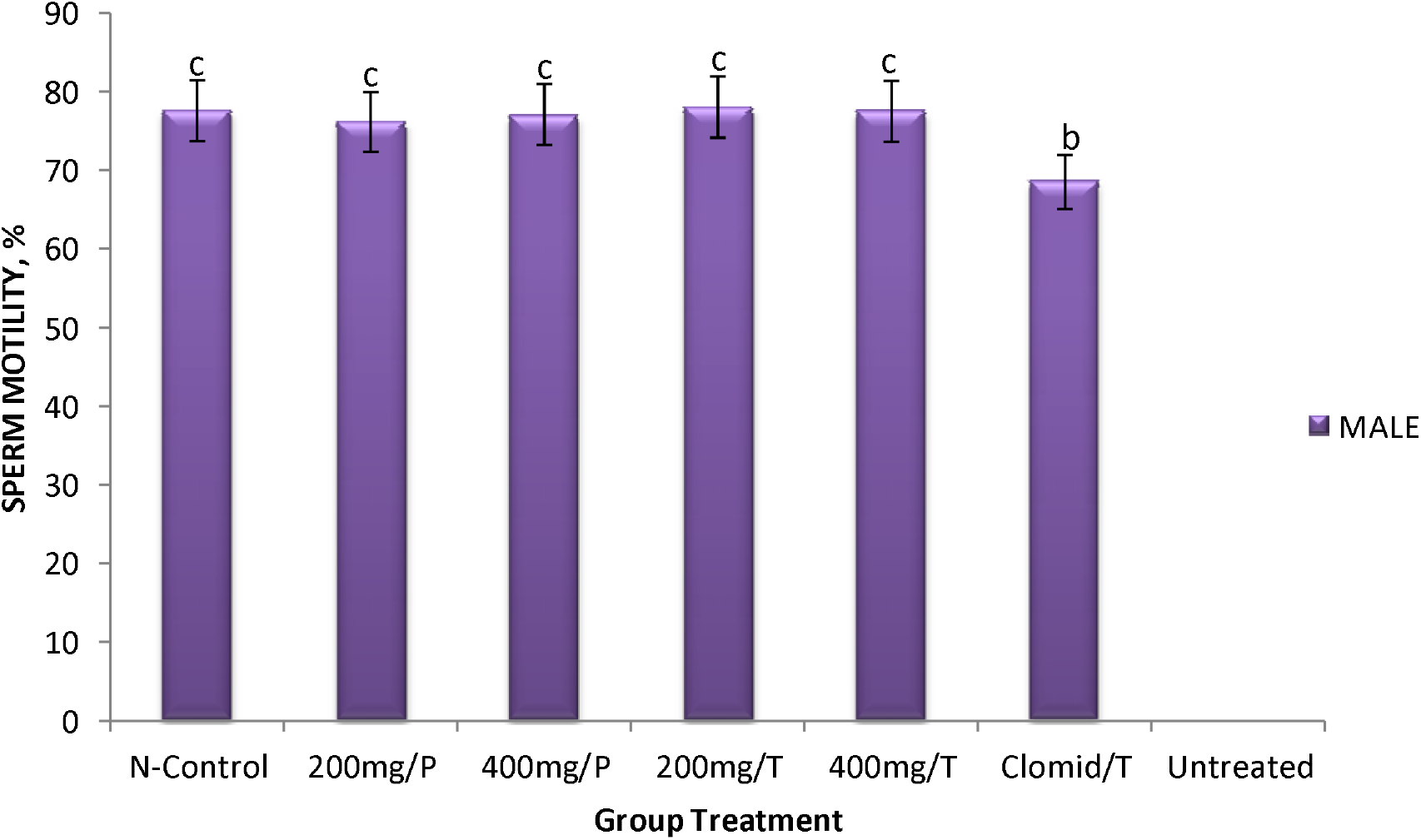
Effect of *Tamarindus indica* fruit pulp extract on sperm motility in male Wistar rats with sodium fluoride induced reproductive dysfunction. Results are Mean ±S.E.M, (n=5). Bars with different letters are significantly different (*p* 0.05).

**FIGURE 8:**
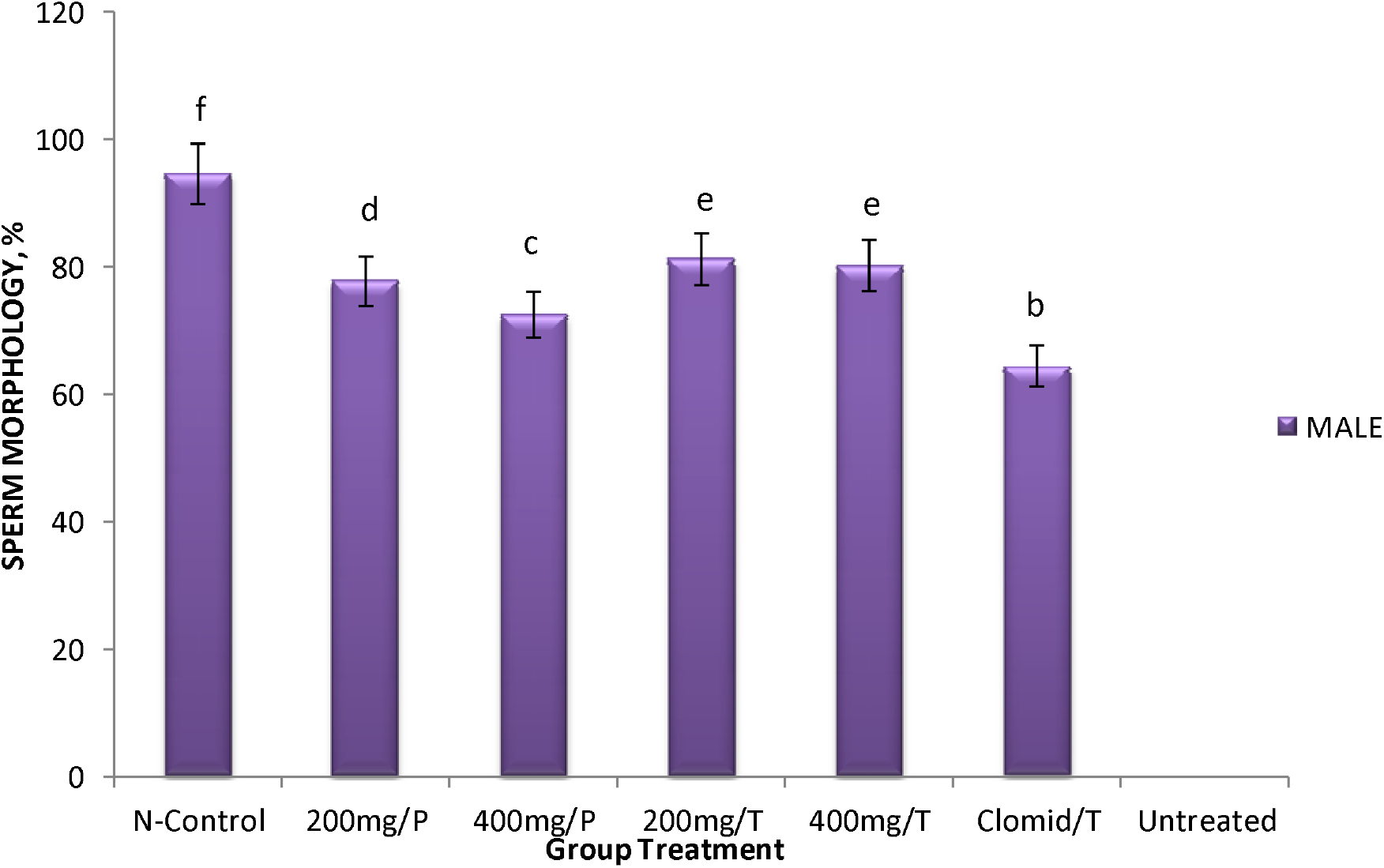
Effect of *Tamarindus indica* fruit pulp extract on sperm morphology in male Wistar rats with sodium fluoride induced reproductive dysfunction. Results are Mean ±S.E.M, (n=5). Bars with different letters are significantly different (*p* 0.05).

**FIGURE 9:**
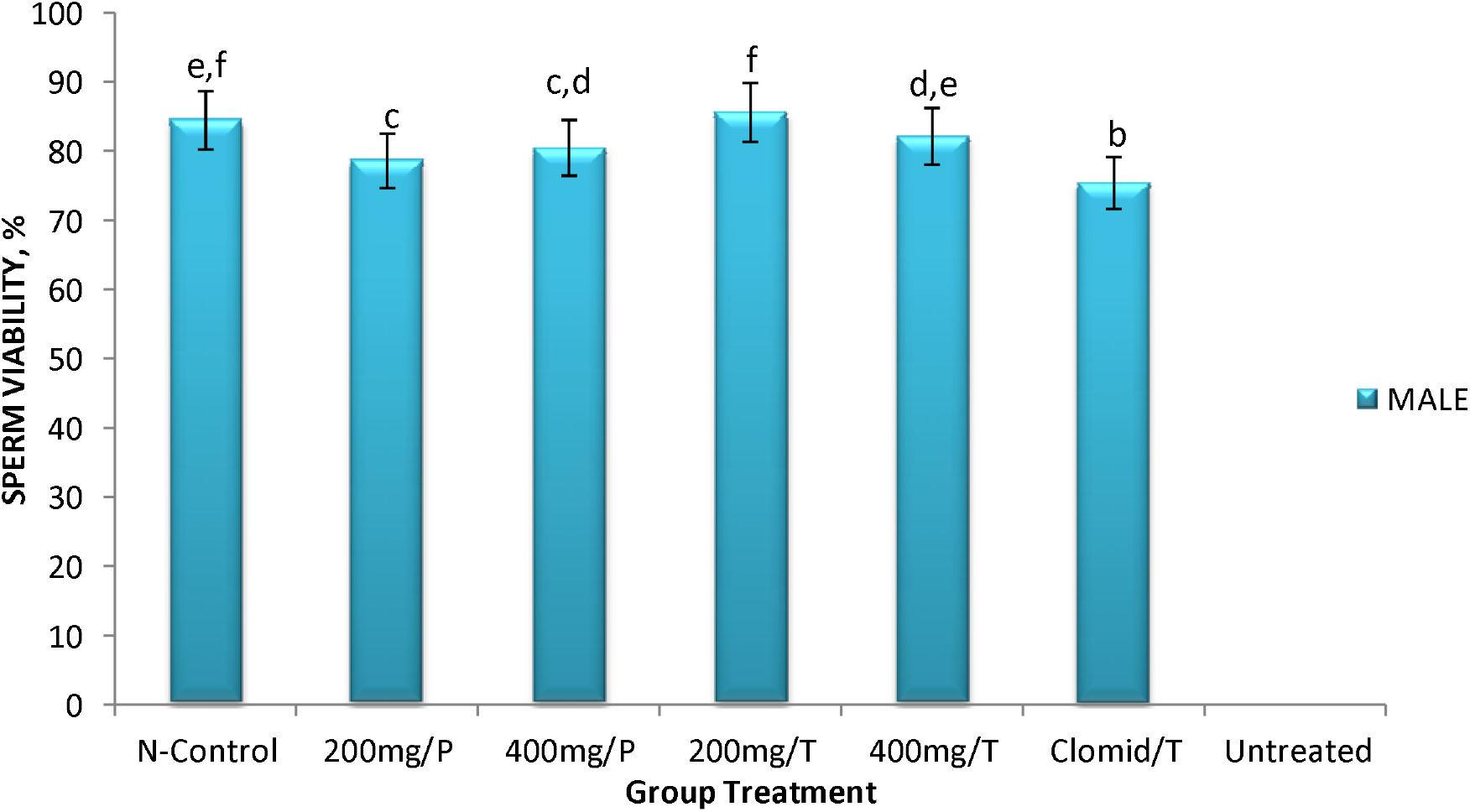
Effect of *Tamarindus indica* fruit pulp extract on sperm viability in male Wistar rats with sodium fluoride induced reproductive dysfunction. Results are Mean ±S.E.M, (n=5). Bars with different letters are significantly different (*p* 0.05).

Plate 1a to g is the representative photomicrograph of the testes of experimental rats after treatment. The control group showed normal testicular histomorphology whereas the model group showed testicular morphology with large interstitial space. The morphology of the rats pre-treated with the extract were comparable with the control group but with comparatively larger interstitial space. The interstitial space of plate of the rats pre-treated with 400 mg/kg dose of the extract showed halo-spaced. The rats treated with the extract after NAF administration also showed normal testicular histomorphology.

## DISCUSSION

Follicle stimulating hormone (FSH) is important for the initiation of spermatogenesis in the Leydig cells and also for the production of testosterone by the Sertoli cells (Ujihara *et al*., 1992). FSH stimulates the theca interna cells for the production of estrogen and also stimulates follicular development prior to ovulation (Durvey and Thakare, 2016). Findings from this study showed that pre-treatment with the extract and administration of the extract after induction of sexual dysfunction raised the serum FSH of the male rats to the pre-treatment value. The result thus suggested that the extract is capable of restoring serum FSH level in reproductive dysfunctional rats similar to the effect of Clomid. This is similar to the findings of Pervez *et al*. (2016) who observed a significant decrease in the serum FSH levels of untreated NaF induced male reproductive dysfunction as compared with the normal control. In this study, 200 mg/kg extract treated (female) group showed statistical elevated FSH level when compared to all other (female) treatment groups. The observed raised FSH level of the female model rats may be attributed to the effect of NaF. Administration of the extract prior to the induction of reproductive dysfunction restored the female FSH level to the control value. This is similar to the observation with Clomid. This could be an indication of an ovulatory disorder, polycystic ovarian syndrome (PCOS) or infertility. This is in accordance with the findings of Durvey and Thakare (2016) who observed an impairment in the oestrus cycle of rats administered with varying concentration of NaF.

Leutenizing hormone stimulates the differentiation of spermatogonia and the production of testost erone by the Sertoli cells (Ujihara *et al*., 1992). Administration of NAF reduced the male LH whereas pre-treatment and post treatment with the extract at both doses raised the LH significantly above the control group. The observed effect could be an indication of a hypergonadotropic effect of *Tamarindus indica*. This result is in consonance with the study of Ma *et al*. (2008) who observed a disturbed hormone levels of each layer of the hypothalamus-hypophysis-testis axis in male rats administered with NaF for a period of time. LH works synergistically with FSH during the development of the follicles and the production of estrogen by the theca interna cells (Ujihara *et al*., 1992). Our observation with the female rats showed that NAF significantly raised the serum LH whereas administration of *Tamarindus indica* extract restored it to the pre-treatment level. The significantly raised LH in the female rats above the control level when pre-treated with 200 mg/kg dose of the *Tamarindus indica* extract may be an indication of the ovulatory status of the rats. This is in consonance with the findings of Durvey and Thakare (2016) who observed an impairment in the oestrus cycle of female rats administered with varying concentration of NaF.

Testosterone is the main androgenic hormone; it is essential for the development of secondary sexual characteristics in males (Bassil *et al*., 2009). It plays an important role in the development of sexual urges. Findings from this study showed a significant reduction in testosterone following administration of NaF. The study reveals that reproductive dysfunction presents a reduction in testosterone level. This result is in accordance with the findings of Pushpalatha *et al*. (2005) who observed a decrease in steroidogenic enzymes in male rats administered with NaF for a period of time. The result is also in accordance with the study of Spittle (2008) who observed a reduced serum testosterone in male rats that received oral administration of NaF for a period of time. This study also reports a significant increase in the testosterone level of the two treatment groups when compared to all the other groups. The extract treated groups performed better than the Clomid treated group. Both groups pre-treated with the extract compared favorably with the normal control group.

Testosterone is the main androgenic hormone; it is essential for the development of secondary sexual characteristics in males (Bassil *et al*., 2009). It plays an important role in the development of sexual urges. This study reports a significant decrease in testosterone level following induction of sexual dysfunction with NAF. Pre-treatment with *Tamarindus indica* prevent this alteration whereas treatment with the extract after induction of sexual dysfunction actually raised the testosterone level above the control value. Our finding agrees with the previous report of Pushpalatha *et al*. (2005) who observed a decrease in steroidogenic enzymes in male rats administered with NaF for a period of time. Spittle (2008) also reported similar findings in male rats that received oral administration of NaF.

Oestrogens are major hormones for growth and development of secondary sexual characteristics in females (Brisken and O’Malley, 2010). It is essential for ovulation, menopause and even pregnancy (Brisken and O’Malley, 2010). Our study showed that sexual dysfunction results in reduction of oestrogen level in rats and that pre-treatment with *Tamarindus indica* extract did not prevent this alteration. Our finding is in agreement with the report of Durvey and Thakare (2016) who observed a decrease in the serum oestrogen levels of female rats administered with NaF. Treatment with the extract at 400 mg/kg dose however boosted the serum oestrogen concentrations as the observed value was above the control value.

Progesterone is a hormone produced majorly by the corpus luteum in the uterus. It is essential to pregnancy, lactation, the ovulatory cycle and parturition (King and Brucker, 2010). We observed in this study that sexual dysfunction elevates serum progesterone level and that pre-treatment with *Tamarindus indica* extract and treatment with the extract after NAF administration restores the progesterone value to the control level. This is in consonance with the findings of Kanwal *et al*. (2016) who observed an impairment in the oestrus cycle in female rats orally administered with NaF in varying concentration (ppm) within a period of 12 days. Pre-treatment with the extract and treatment after NaF administration did not restore the progesterone level to the pre-treatment level

The result of the sperm count, motility, morphology and viability of the male experimental rats showed a reduction of these parameters in the model group signifying that sexual dysfunction lowers sperm count, motility, morphology and viability. These alterations in sperm parameters may be attributable to the decrease in steroidogenic enzymes caused by oral exposure to the NaF. This finding is supported by the report of Pushpalatha *et al*. (2005). The sperm progressivity across all the treatment groups was also observed to be a good, forward, directional movement except for the model and the positive control group where the progressivity was observed to be poor. The highest sperm count was observed in the rats pre-treated with 400 mg/kg of *Tamarindus indica* extract while no motility was observed in the model group. This observation is in conformity with; the findings of Pushpalatha *et al*. (2005) and Spittle (2008) who both reported a decrease in sperm motility in male rats that received oral administration of NaF.

The result of the histopathological analysis showed that the oral administration of NaF over a certain period results in large, expanded, halo spaced instertitium and it is observed that the pre-treatment with *Tamarindus indica* fruit pulp extract has no significant effect in contracting the expanded interstitial space caused by the oral administration of NaF. However, when administered after the induction of sexual dysfunction, *Tamarindus indica* showed a positive effect in contracting the expanded interstitium caused by the oral administration of NaF. This result is in accordance with the findings of Narayana and Chinoy (1994) and Wan *et al*. (2006b) who both observed a decrease in the diameter of seminiferous tubules and a significant change in the diameter of Leydig cells in male rats that have received oral administration of NaF respectively.

## CONCLUSION

Our study revealed administration of the fruit pulp extract of *Tamarindus indica* have both prophylactic and ameliorative effect on sodium fluoride induced reproductive dysfunctional albino rats. This is evidenced by the hormonal, histopathology and sperm function indices findings reported in this study. Our findings showed that this effect is dose dependent thus suggesting that *Tamarindus indica* may be exploited as an alternative to conventional drugs and methods in the management of infertility.

Figure 7 represents the treatment effect on sperm motility. There was no statistical difference between the control group and all the extract treated groups. The lowest sperm motility value was observed in the modal group.

### 4.4 EFFECT OF TREATMENT ON TESTIS HISTOLOGY

**FIGURE 12.**
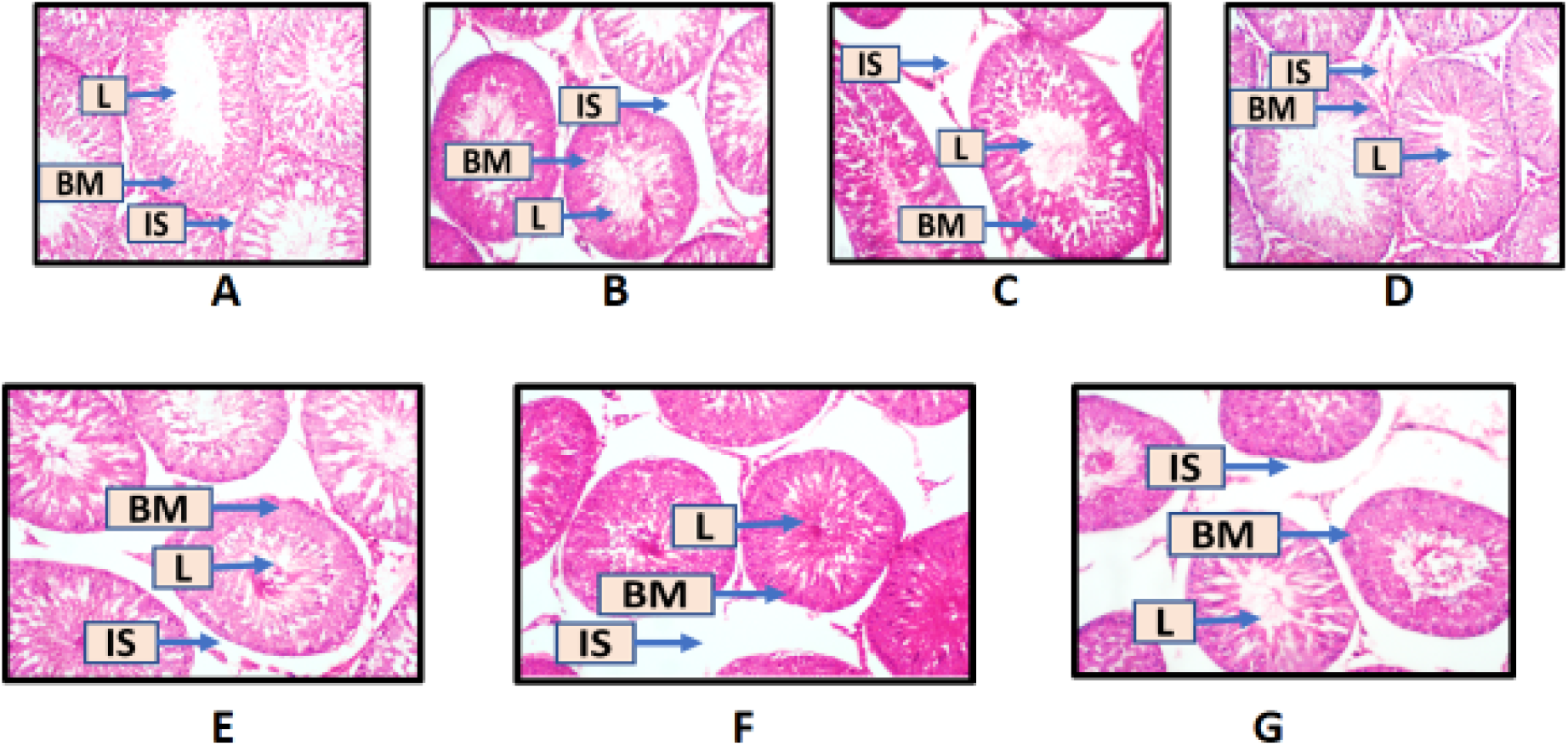
REPRESENTATIVE PHOTOMICROGRAPH OF THE TESTES AFTER TREATMENT. **A =** the representative photomicrograph of the testes of male experimental rats in the normal control group; **A** presents with normal testicular histomorphology. **B =** Representative photomicrograph of the testes of male experimental rats in the 200 mg/kg preventive group, showing a high-power magnification (x100 mag) of the seminiferous tubules. (L= Lumen, BM= Basement membrane, IS= Interstitial space). **C =** Representative photomicrograph of the testes of male experimental rats in the 400 mg/kg preventive group, showing a high-power magnification (x100 mag) of the seminiferous tubules. (L= Lumen, BM= Basement membrane, IS= Interstitial space). **B** and **C** has a normal histomorphology with comparatively larger interstitial space; the interstitial space of **B** is halo-spaced. **D =** Representative photomicrograph of the testes of male experimental rats in the 200 mg/kg treatment group, showing a high-power magnification (x100 mag) of the seminiferous tubules. (L= Lumen, BM= Basement membrane, IS= Interstitial space). **E =** Representative photomicrograph of the testes of male experimental rats in the 400 mg/kg treatment group, showing a high-power magnification (x100 mag) of the seminiferous tubules. (L= Lumen, BM= Basement membrane, IS= Interstitial space). **D** and **E** presents with normal testicular histomorphology. **F =** Representative photomicrograph of the testes of male experimental rats in the positive control group, showing a high-power magnification (x100 mag) of the seminiferous tubules. (L= Lumen, BM= Basement membrane, IS= Interstitial space). **G =** Representative photomicrograph of the testes of male experimental rats in the negative control group, showing a high-power magnification (x100 mag) of the seminiferous tubules. (L= Lumen, BM= Basement membrane, IS= Interstitial space). **F** presents with relatively expanded interstitial while **G** presents with large interstitial space.

## Notes

### Competing Interest Statement

The authors have declared no competing interest.

